# Taxonbridge: an R package to create custom taxonomies based on the NCBI and GBIF taxonomies

**DOI:** 10.1101/2022.05.02.490269

**Authors:** Werner P. Veldsman, Giulia Campli, Sagane Dind, Valentine Rech de Laval, Harriet B. Drage, Robert M. Waterhouse, Marc Robinson-Rechavi

## Abstract

**Summary:** Biological taxonomies establish conventions by which researchers can catalogue and systematically compare their work using nomenclature such as species binomial names and reference identifiers. The ideal taxonomy is unambiguous and exhaustive; however, no such single taxonomy exists, partly due to continuous changes and contributions made to existing taxonomies. The degree to which a taxonomy is useful furthermore depends on context provided by such variables as the taxonomic neighbourhood of a species (e.g., selecting arthropod or vertebrate species) or the geological time frame of the study (e.g., selecting extinct *versus* extant species). Collating the most relevant taxonomic information from multiple taxonomies is hampered by arbitrarily defined identifiers, ambiguity in scientific names, as well as duplicated and erroneous entries. The goal of *taxonbridge* is to provide tools for merging the Global Biodiversity Information Facility (GBIF) Backbone Taxonomy and the United States National Center for Biotechnology Information (NCBI) Taxonomy in order to create consistent, deduplicated and disambiguated custom taxonomies that reference both extant and extinct species.

**Availability:** *Taxonbridge* is available as a package in the Comprehensive *R* Archive Network (CRAN) repository: https://CRAN.R-project.org/package=taxonbridge.

## INTRODUCTION

A comprehensive map of the current understanding of the evolutionary relationships amongst organisms is a prerequisite cornerstone of most downstream comparative analyses. However, since there is a plurality of taxonomic databases with heterogeneous specification and scope, it is often challenging to collate an authoritative taxonomy in a consistent manner. This is especially true for large studies with hundreds to thousands of organisms spanning disparate taxa and geological time frames. Efforts to develop authoritative references have led to several key taxonomy resources such as the Catalogue of Life (COL) (Catalogue of Life, 2022), the Global Biodiversity Information Facility (GBIF) Backbone Taxonomy (GBIF: The Global Biodiversity Information Facility, 2022), the Open Tree of Life (OToL) reference taxonomy (Rees and Cranston, 2017), or the United States National Center for Biotechnology Information (NCBI) Taxonomy (Schoch *et al*. 2020). With different strategies and priorities, these and other more specialised resources serve the needs of different yet often overlapping research communities. For example, the NCBI Taxonomy includes organism names and classifications for species with sequences from the International Nucleotide Sequence Database Collaboration, meaning that species without sequence data such as most extinct species are underrepresented. In contrast, the GBIF Backbone Taxonomy references more extensive collections of extinct and extant species from at least 100 source taxonomies, including the COL, allowing the GBIF to integrate name-based information from different resources. Factors that confound taxonomic integration also include arbitrarily assigned identifiers, ambiguity in binomial naming, as well as duplicated and erroneous entries (Zizka et al., 2020). Despite progress in harmonisation and interoperability efforts, for example, in terms of data processing (GBIF Secretariat, 2021) as well as in the conceptualisation of integration strategies (Waterhouse *et al*., 2021), the GBIF and NCBI taxonomies are currently not readily integrated. And while the OToL reference taxonomy (v3.3 generated on 1 June 2021) incorporates GBIF and NCBI taxonomic data, their taxonomic data does not include lineage data per taxonomic source, which hampers comparison of lineage information to detect ambiguity. We thus present *taxonbridge*, an *R* package with generic tools that allow users to generate near real-time custom merged GBIF and NCBI taxonomies that are consistent, deduplicated and disambiguated and that reference both extant and extinct species.

## IMPLEMENTATION

*Taxonbridge* is implemented using a functional programming approach with S3 class constructs. Source code was written using *R base* v4.1.2 and *devtools* v2.4.3. *Taxonbridge* consists of seventeen methods grouped into four categories or “steps” (Figure 1).

**Figure 1:**
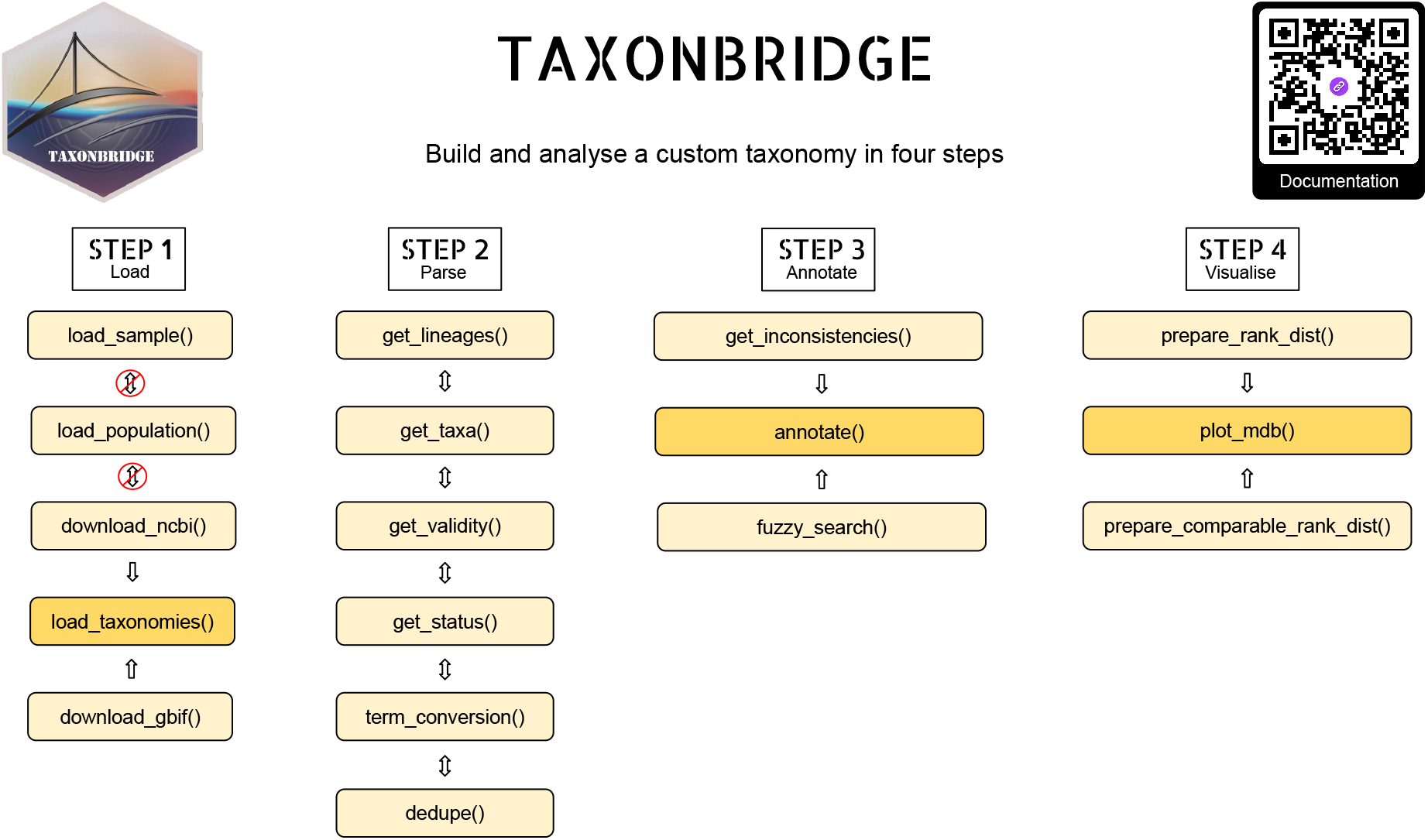
Workflow. The *taxonbridge* workflow consists of seventeen methods grouped into four categories. The four categories (“steps”) are designed to be used consecutively and are respectively useful for parsing, filtering, annotation, and visualisation of taxonomic data.

The first step consists of five methods that allow loading and parsing of GBIF and NCBI taxonomic data. Parsing of the NCBI taxonomy is supported by the external software library *Taxonkit* v.0.9.0 (Shen and Ren, 2021). The remainder of imported third-party packages are all *R* packages themselves and include *dplyr* v1.0.8, *ggplot2* v3.3.5, *purrr* v0.3.4, *rje* v1.10.16, *stringr* v1.4.0, *vroom* v1.5.7, and *withr* v2.5.0. NCBI and GBIF taxonomies can be downloaded, parsed and merged using a single method or multiple methods if needed. The second step consists of six methods that allow filtering by taxa and status, retrieval of lineage data, validation based on lineage data, conversion of synonymous terms, and deduplication. The six methods in the second step can be used repetitively and in any order. The third and fourth steps each consist of three methods, two of which generate data that should be passed to the third (see Figure 1). The purpose of the third and fourth steps are to allow annotation and visualisation of a custom taxonomy. Example use cases for *taxonbridge* are provided in a vignette that has been made available at the CRAN repository webpage.

## DEMONSTRATION

Consider a study in which a researcher who is in the process of collating a taxonomy would like to determine which species included in the study have binomial names that are identical to the binomial names of species that are not of interest to the study. For example, a researcher may wish to determine which arthropods have identical binomial names to species that are assigned to both different families and different kingdoms, bearing in mind that a difference may arise from either ambiguity (e.g. two different species sharing a common name) or inconsistent naming (e.g. different nomenclature used to refer to the same species). An exercise to answer this question was carried out on 12 April 2022. *Taxonbridge* enabled retrieval of the latest available taxonomic information, downloading, parsing and merging the GBIF backbone taxonomy and the NCBI taxonomy in less than five minutes. In the GBIF backbone taxonomy 6,186,255 entries out of 6,957,235 contained scientific names (including uninomials, binomials and other matching phrases), while in the NCBI taxonomy all 2,414,305 entries contained scientific names. A total of 695,220 matches between the GBIF and NCBI entries could be made based on scientific names. The resultant taxonomy contained a total of 3,177,308 entries that were assigned to the phylum Arthropoda, of which 231,922 had lineage data in both the GBIF and NCBI. At the family level, *taxonbridge* detected 26,900 arthropod entries with a discrepancy between the GBIF and NCBI lineage data, and, at the kingdom level, found 4,567 arthropod entries with a discrepancy between the GBIF and NCBI lineage data. The intersection of these two non-corresponding sets revealed 96 binomial names that differ at both the kingdom and family levels. For comparison, only 37 of these ambiguous or inconsistent names were present as double entries in the OToL reference taxonomy v3.3, while 47 names were present as single entries, and 12 names were absent. We attempted to reproduce and update the OToL reference taxonomy but encountered multiple issues during installation. An interesting example from the group of 96 ambiguous or inconsistent names that we detected using Taxonbridge is that of *Drosophila badia*, which is the assigned name of a vinegar fly (Animalia: Drosophilidae) in the GBIF backbone taxonomy but the assigned name of a mushroom (Fungi: Psathyrellaceae) in the NCBI taxonomy. Curiously, there is a mushroom from the Psathyrellaceae family that is assigned the name *Psathyrella niveobadia* in both the GBIF and NCBI taxonomy, with *Drosophila niveobadia* recognised as a synonym by the GBIF and as a basionym by the NCBI. This example highlights the need for taxonomic curation in biological taxonomies, and the efficacy of *taxonbridge* in promoting the discovery of ambiguity and inconsistencies in biological taxonomies.

## ACKNOWLEDGEMENTS

The authors acknowledge the contributions of all members of the arthropod moulting project: Allison Daley, Sinéad Lynch, Kenneth Kim, Olga Volovych, Jonathan Antcliffe, Idan Sheizaf, Asya Novikov and Ariel D. Chipman. *Taxonbridge* was developed with funding from a Swiss National Science Foundation (SNSF) Sinergia award [grant number 198691].

## Notes

https://CRAN.R-project.org/package=taxonbridge

https://github.com/MoultDB/taxonbridge

### Competing Interest Statement

The authors have declared no competing interest.

